# Serum metabolomics and incidence of atrial fibrillation: the Atherosclerosis Risk in Communities (ARIC) Study

**DOI:** 10.1101/368118

**Authors:** Alvaro Alonso, Bing Yu, Yan V. Sun, Lin Y. Chen, Laura R. Loehr, Wesley T. O’Neal, Elsayed Z. Soliman, Eric Boerwinkle

## Abstract

We have previously identified associations of two circulating secondary bile acids (glycocholenate and glycolithocolate sulfate) with atrial fibrillation (AF) risk among blacks. We aimed to replicate these findings in an independent sample including both whites and blacks, and performed a new metabolomic analysis in the combined sample. We studied 3,922 participants from the ARIC cohort followed between 1987 and 2013. Of these, 1,919 had been included in the prior analysis and 2,003 were new samples. Metabolomic profiling was done in baseline serum samples using gas and liquid chromatography mass spectrometry. AF was ascertained from electrocardiograms, hospitalizations, and death certificates. We used multivariable Cox regression to estimate hazard ratios (HR) and 95% confidence intervals (95%CI) of AF by one standard deviation difference of metabolite levels. Over a mean follow-up of 20 years, 608 participants developed AF. Glycocholenate sulfate was associated with AF in the replication and combined samples (HR 1.10, 95%CI 1.00, 1.21 and HR 1.13, 95%CI 1.04, 1.22, respectively). Glycolithocolate sulfate was not related to AF risk in the replication sample (HR 1.02, 95%CI 0.92, 1.13). An analysis of 245 metabolites in the combined cohort identified three additional metabolites associated with AF after multiple-comparison correction: pseudouridine (HR 1.18, 95%CI 1.10, 1.28), uridine (HR 0.86, 95%CI 0.79, 0.93) and acisoga (HR 1.17, 95%CI 1.09, 1.26). To conclude, we replicated a prospective association between a previously identified secondary bile acid, glycocholenate sulfate, and AF incidence, and identified new metabolites involved in nucleoside and polyamine metabolism as markers of AF risk.

## INTRODUCTION

Atrial fibrillation (AF), a common cardiac arrhythmia, is a major risk factor for stroke and other cardiovascular diseases.^1^ Application of metabolomics, the systematic investigation of all small molecules in a biological system, to the study of AF risk could deepen our understanding of AF pathogenic pathways as well as contribute to the discovery of novel disease biomarkers.^2^ To date, however, metabolomic studies in this area have been few and limited in sample size. In an analysis of metabolomic data from 1,919 black participants in the community-based Atherosclerosis Risk in Communities (ARIC) study, including 183 who were newly diagnosed with AF, we reported an association of higher circulating levels of two secondary bile acids, glycolithocholate sulfate and glycocholenate sulfate, with incidence of AF, but no replication in independent cohorts was available.^3^ More recently, a report from the mostly European-American Framingham Heart Study including 2,458 participants with targeted metabolomic profiling, of which 156 developed AF, did not identify any molecule significantly associated with AF incidence after adjustment for multiple comparisons.^4^ Additional studies are required to replicate previous findings and increase statistical power for novel discoveries.

In this manuscript, as a follow-up to our previous study in the ARIC cohort, we extend the metabolomic assessment to 2,003 additional ARIC participants. We aimed to replicate the findings from the prior ARIC analysis in the additional ARIC participants and to conduct a new hypothesis-generating analysis in the combined sample of 3,922 participants.

## METHODS

### Study population

In 1987-89, the ARIC study examined 15,792 men and women 45-64 years of age recruited from four communities in the United States (Forsyth County, NC; Jackson, MS; Minneapolis suburbs, MN; Washington County, MD).^5^ Participants were mostly white in the Minneapolis and Washington County sites, white and black in Forsyth County, while only black were recruited in Jackson. After their baseline exam, participants underwent follow-up visits in 1990-92, 1993-95, 1996-98, 2011-13, and 2016-17. Participants have been followed up via annual phone calls (semiannual since 2012). For the current analysis, we included 3,922 participants with available metabolomic data and without evidence of AF at baseline. The ARIC study has been approved by institutional review boards at all participating institutions (University of Minnesota, University of North Carolina, University of Mississippi Medical Center, Johns Hopkins University, and University of Texas Health Sciences Center). Participants provided written informed consent at baseline and follow-up visits. This research was conducted in accordance with relevant guidelines and regulations pertaining to human subjects research in the United States.

### Metabolomic profiling

As previously described, 1,977 randomly selected blacks in the Jackson field center had serum metabolomic profiling performed in 2010 in samples obtained at study baseline in 1987-89.^6^ The samples had been stored at −80°C and were assayed with an untargeted, gas chromatography/mass spectrometry and liquid chromatography/mass spectrometry-based metabolomic protocol by Metabolon, Inc. (Durham, NC). Similarly in 2014, serum samples from an additional 2,055 randomly selected participants (76% white, 24% black) collected in 1987-89 and stored since then at −80°C were assayed by Metabolon, Inc. using the same protocol. Brief methodological details are provided here. Sample preparation begins with four different fractionation steps to extract all polar and nonpolar molecules with a mass of 50–1,500 daltons. Once these small molecules have been separated out through these four extraction steps, they are pooled back together, and that sample is then split for analysis by two different platforms—a liquid chromatography–mass spectrometry system (LC-MS) and a gas chromatography–mass spectrometry system (GC-MS). Both of these platforms are used because small molecules can be very polar as well as very nonpolar; the two chromatography methods work well together for profiling of most of the small molecules in the samples. In house software is then used to identify all of the ions that are scanned by the spectrometers. Using automated processing techniques based on the biological variation of the compounds within samples, the researchers are able to reconstruct the original molecules to which the ions belonged before going through the system. With the help of a standard chemical library, the molecules are identified, and their amounts are quantified. The end result is a data set that identifies all the small molecules seen in the sample and their relative amounts.

We selected a set of 97 samples to measure their metabolome profiles using baseline serum samples at both 2010 and 2014. We calculated the Pearson correlation coefficients (r) between the 97 pairs for shared metabolites. For the present study, we limited the analysis to metabolites with: 1) no more than 25% missing values, and 2) Pearson correlation coefficients ≥ 0.3 between 2010 and 2014 measurements. After applying these criteria, 245 named metabolites were included. To evaluate the stability of samples in long-term storage, we compared metabolomic measures done at 2014 and 2016 with standard clinical laboratory measures done at ARIC baseline (1989) for urea, glucose, and cholesterol. All three metabolites showed Pearson correlation coefficients ≥0.65.

### Ascertainment of atrial fibrillation

We have described elsewhere the details about AF ascertainment in the ARIC cohort.^7^ Briefly, we identified AF cases through the end of 2013 from three sources: electrocardiograms (ECG) done at scheduled study visits, discharge diagnosis codes from hospitalizations, and death certificates. At all study visits, participants underwent a standard 12-lead 10-second ECG, which was transmitted electronically to the ARIC ECG reading center at EPICARE (Wake Forest School of Medicine, Winston-Salem, NC) for review and analysis using the GE Marquette 12-SL program (GE Marquette, Milwaukee, WI). A computer algorithm identified the presence of AF in the ECG, with a cardiologist confirming the diagnosis.

Participants’ hospitalizations during follow-up were identified through phone calls and surveillance of local hospitals. Trained abstractors collected information from these hospitalizations, including all discharge codes. We considered AF present if ICD-9-CM codes 427.31 or 427.32 were listed as discharge diagnoses in any given hospitalization. We excluded AF cases associated with open cardiac surgery. We and others have demonstrated adequate validity of this approach for the ascertainment of AF.^7,8^ Finally, we also defined AF from death certificates if ICD-9 427.3 or ICD-10 I48 were listed as any cause of death.

### Covariates

During the baseline visit, participants self-reported age, sex, race, and smoking history and underwent a physical exam that included measurements of blood pressure, weight, and height. Blood glucose and lipid concentrations were measured using standard methods in baseline samples. Estimated glomerular filtration rate (eGFR) was calculated from serum creatinine using the CKD-EPI equation.^9^ Diabetes was defined if the participant had fasting blood glucose ≥126 mg/dL, non-fasting blood glucose >200 mg/dL, used antidiabetic medications, or reported a physician-diagnosis of diabetes. Prevalent heart failure was defined according to Gothenburg criteria,^10^ while prevalent coronary heart disease was based on self-reported information. Also at baseline, participants underwent a standard 12-lead 10-second electrocardiogram, which was processed at EPICARE (Wake Forest School of Medicine, Winston-Salem, NC). PR duration, P wave axis and P wave terminal force in V1 were all automatically measured. Abnormal P wave axis was defined as any P wave axis value outside 0 to 75 degrees, while elevated P wave terminal force in V1 was defined if P wave terminal force was >4,000 μV*ms. Genome-wide and exome genotyping of ARIC participants has been done using the Affymetrix 6.0 and the Illumina HumanExome Beadchip v1.0, as described elsewhere.^11^

### Statistical analysis

We conducted two separate sets of analyses. In the first one, we aimed to replicate the findings from our prior ARIC publication, estimating the association of glycolithocholate sulfate and glycocholenate sulfate with AF incidence in 2,003 participants without AF at baseline not included in our published analysis. A second analysis combined participants from the two metabolomic assessment batches (n = 3,922). We used a modified Bonferroni correction to determine statistical significance.^12^ Using this approach, p-values less than 3.538 × 10^−4^ were considered statistically significant for 245 tested metabolites. Additionally, since the Bonferroni correction can be too conservative and miss true associations, we alternatively defined statistical significance using a false discovery rate (FDR) of 5% applying the Benjamini-Hochberg procedure.^13^

For all analyses, the association of individual metabolites with the incidence of AF was estimated with Cox proportional hazards regression. Time of follow-up was defined as the time in days from the baseline visit to incidence of AF, death, loss to follow-up or December 31, 2013, whichever occurred earlier. Metabolites were mean centered and modeled as continuous variables in standard deviation units. Missing values were imputed with the lowest detected value in each batch. We ran three separate models with increasing number of covariates. A first model adjusted for age, sex, race, center, and batch (when applicable). A second model additionally adjusted for smoking, body mass index, systolic blood pressure, hypertension medications, diabetes mellitus, history of heart failure, and history of coronary heart disease. A final model additionally adjusted for eGFR. We selected model covariates based on prior knowledge of risk factors for AF.^14^ We assessed effect measure modification by race and sex using stratified analysis. The dose-response shape of the association between metabolite concentration and AF incidence was evaluated modeling metabolites using a restricted cubic spline with five knots. To test the robustness of the observed significant associations, we conducted a series of sensitivity analyses, adjusting for blood lipids and lipid-lowering medications and excluding participants with a prior history of prevalent coronary heart disease or heart failure, as well as adjusting for aspartate aminotransferase (AST) and alanine aminotransferase (ALT), measured in visit 2 samples, in the analyses of bile acids.

We conducted several additional analyses to explore potential mechanisms of the association between metabolites and AF incidence. First, we evaluated the association of statistically significant metabolites with electrocardiographic endophenotypes of AF risk using linear regression (PR duration, in ms) or logistic regression (abnormal P wave axis and elevated P wave terminal force in V1). Second, we evaluated the association of statistically significant metabolites with 23 single nucleotide polymorphisms (SNPs) associated with AF in a prior genome-wide association study (GWAS) from the AFGen consortium, and a genetic score calculated by adding the number of risk alleles weighted by the beta coefficient from the published genome-wide study.^11^ Finally, we explored whether variation in rs2272996 in gene *VNN1*, a SNP previously related to circulating concentrations of acisoga (one of the metabolites associated with AF incidence in this analysis),^15^ was associated with AF incidence in the latest GWAS of AF.

## RESULTS

Of 15,792 participants in the ARIC cohort, the present analysis included 3,922 with available metabolomic data and free of AF at baseline, 1,919 of them included in our previous publication and 2,003 with newly available data. Participants were followed up for a mean (standard deviation) of 20.4 (7.0) years, during which 608 AF events were identified (incidence rate, 7.6 cases per 1,000 person-years). Table 1 reports participants’ characteristics overall and by AF incidence status during follow-up. As expected, participants who developed AF during follow-up were older, had higher systolic blood pressure and worse kidney function at baseline. They were also more likely to be white, male and have a baseline diagnosis of diabetes, heart failure or coronary heart disease.

**Table 1.** Selected baseline characteristics by atrial fibrillation (AF) status during follow-up in 3,922 participants with available metabolomic data and free of AF at baseline, ARIC study, 1987-89

| Baseline | Overall | No incident AF | Incident AF |
| --- | --- | --- | --- |
| N | 3,922 | 3,314 | 608 |
| Age, years | 54 (6) | 53 (6) | 56 (6) |
| Women, % | 60 | 62 | 50 |
| Race |  |  |  |
| Black, % | 61 | 64 | 47 |
| White, % | 39 | 36 | 53 |
| Body mass index, kg/m<sup>2</sup> | 29 (6) | 29 (6) | 30 (6) |
| Current smoker, % | 28 | 27 | 29 |
| Systolic blood pressure, mmHg | 125 (21) | 124 (21) | 129 (22) |
| Anti-hypertensive medication, % | 32 | 31 | 39 |
| Diabetes, % | 14 | 13 | 19 |
| eGFR, mL/min/1.73 m<sup>2</sup> | 99 (18) | 100 (18) | 94 (19) |
| Prevalent heart failure, % | 5.1 | 4.5 | 8.4 |
| Prevalent coronary heart disease, % | 4.8 | 4.0 | 9.2 |
| Values correspond to mean (standard deviation) or percentages. eGFR: estimated glomerular filtration rate |  |  |  |

In an initial analysis, we aimed to replicate the findings from our previous publication showing that higher levels of glycolithocholate sulfate and glycocholenate sulfate were associated with increased risk of AF. In an age and sex-adjusted analysis including 2,003 participants and 386 incident AF events, higher levels of glycocholenate sulfate but not of glycolithocholate sulfate were associated with AF incidence in the replication analysis (Table 2, Model 1). The association of glycocholenate sulfate with incidence of AF became weaker after multivariable adjustment (HR 1.10, 95%CI 1.00, 1.21 per 1-SD difference; Table 2, Model 2). Given the strong attenuation after multivariable adjustment, we explored if any individual covariate was responsible for this change. Adding each covariate to Model 1 individually did not point to any particular variable as responsible for the attenuation (Supplementary Figure 1). The hazard ratio (HR) and 95% confidence interval (CI) of AF per 1-standard deviation (SD) difference in glycocholenate sulfate in the combined derivation and replication samples was 1.23 (95%CI 1.14-1.32, p = 9.5 × 10^−8^) in minimally adjusted models and 1.13 (95%CI 1.04, 1.22, p = 0.003) after additional adjustment for cardiovascular risk factors. Additional adjustment for concentrations of ALT and AST in 3,401 participants with available information on liver enzymes did not modify the associations (HR 1.15, 95%CI 1.07, 1.23, p = 2.5 × 10^−5^). Analysis stratified by race and sex showed a weaker association between glycolithocholate sulfate and AF in whites compared to blacks (HR 1.04, 95%CI 0.94, 1.16 versus HR 1.19, 95%CI 1.10, 1.28, p for interaction = 0.05). No other interactions were identified (Supplementary Figures 2 and 3).

**Table 2.** Association of two secondary bile acids (glycocholenate sulfate and glycolithocholate sulfate) with incidence of AF, by analytical batch. Hazard ratios per 1-standard deviation difference. ARIC study, 1987-2013

|  | First batch<br>(N = 1919; AF = 222) |  | Second batch<br>(N = 2003; AF = 386) |  | Combined sample<br>(N = 3,922; AF = 608) |  |
| --- | --- | --- | --- | --- | --- | --- |
|  | HR (95%CI) | p-value | HR (95%CI) | p-value | HR (95%CI) | p-value |
| Glycocholate sulfate |  |  |  |  |  |  |
| Model 1 | 1.27 (1.16, 1.39) | $1.9 \times 10^{-7}$ | 1.21 (1.10, 1.33) | 0.0001 | 1.23 (1.14, 1.32) | $9.5 \times 10^{-8}$ |
| Model 2 | 1.20 (1.08, 1.33) | 0.0006 | 1.10 (1.00, 1.21) | 0.05 | 1.13 (1.04, 1.22) | 0.003 |
| Glycolithocholate sulfate |  |  |  |  |  |  |
| Model 1 | 1.22 (1.13, 1.31) | $1.4 \times 10^{-7}$ | 1.02 (0.93, 1.13) | 0.69 | 1.09 (1.01, 1.17) | 0.03 |
| Model 2 | 1.21 (1.11, 1.31) | $5.5 \times 10^{-6}$ | 1.02 (0.92, 1.13) | 0.67 | 1.07 (0.99, 1.15) | 0.11 |
| Model 1 adjusted for age, sex and race, center and batch where applicable. Model 2 additionally adjusted for smoking, body mass index, systolic blood pressure, use of antihypertensive medication, diabetes, prevalent heart failure, and prevalent coronary heart disease. |  |  |  |  |  |  |

Subsequently, we performed a metabolome-wide, hypothesis-free analysis combining the two study samples. Of the 245 studied metabolites, 9 were associated with the incidence of AF with p-values <0.001 after multivariable adjustment and 11 had FDR-adjusted p-value<0.05 (Table 3, Model 2). These metabolites included molecules involved in the metabolism of pyrimidines (pseudouridine and uridine), polyamines (acisoga), amino acids (N-acetylalanine and N-acetylthreonine), dipeptides (gamma-glutamylisoleucine and gamma-glutamylleucine), and bile acids (glycoursodeoxycholate and glycochenodeoxycholate), as well as one lysolipid (1-docosahexaenoylglycerophosphocholine), and a xenobiotic (O-sulfo-L-tyrosine). Pearson correlation coefficients for these metabolites between repeated measures in 97 samples as well as percentage of observations with missing values are presented in Supplementary Table 1. Three of these molecules, pseudouridine, acisoga, and uridine, were significantly associated with AF with p-values < 3.538 × 10^−4^. Specifically, higher levels of pseudouridine and acisoga were associated with higher rates of AF (HR 1.18, 95%CI 1.10, 1.28 and 1.17, 95%CI 1.09, 1.26, respectively) while higher uridine levels were associated with reduced AF rates (HR 0.86, 95%CI 0.79, 0.93). Complete results for the 245 metabolites are available as a supplementary file. The correlation matrix of the 11 metabolites is shown in Supplementary Table 2. Uridine was not correlated with pseudouridine (r = −0.02) or acisoga (r = −0.03), though there was a modest association between pseudouridine and acisoga (r = 0.42). Associations for pseudouridine and acisoga weakened, but were still present, in a model including the 3 metabolites simultaneously (HR 1.16, 95%CI 1.06, 1.26 for pseudouridine, HR 1.11, 95%CI 1.02, 1.20 for acisoga). The inverse association between uridine and AF risk did not change after adjustment for pseudouridine and acisoga (HR 0.85, 95%CI 0.79, 0.92). The association remained essentially unchanged after adjustment for blood lipids and in those without CVD (Supplementary Table 3). Figure 1 presents the dose-response associations of pseudouridine, acisoga, and uridine with AF risk, which were approximately linear for the three molecules. Multivariable adjustment led to meaningful attenuation in the association of pseudouridine with AF. None of the individual covariates in the multivariable model seemed particularly responsible for this attenuation, as evaluated by adding each covariate individually to the minimally adjusted model (Supplementary Figure 1). Associations were similar across race and sex groups (Supplementary Figures 2 and 3).

**Table 3.** Association of individual metabolites with incidence of atrial fibrillation, ARIC study, 1987–2013. Hazard ratios per 1-standard deviation difference. Only metabolites with an FDR-adjusted p-value <0.05 in the multivariable model 2 are shown.

| Metabolite | Model 1 |  |  | Model 2 |  |  | Model 3 |  |  |
| --- | --- | --- | --- | --- | --- | --- | --- | --- | --- |
|  | HR (95%CI) | p-value | FDR p | HR (95%CI) | p-value | FDR p | HR (95%CI) | p-value | FDR p |
| Pseudouridine | 1.31 (1.22, 1.41) | 4.5x10<sup>-13</sup> | 5.5x10<sup>-11</sup> | 1.18 (1.10, 1.28) | 1.7x10<sup>-5</sup> | 4.0x10<sup>-3</sup> | 1.16 (1.06, 1.27) | 9.6x10<sup>-4</sup> | 0.031 |
| Acisoga | 1.20 (1.12, 1.30) | 1.3x10<sup>-6</sup> | 2.3x10<sup>-5</sup> | 1.17 (1.09, 1.26) | 4.0x10<sup>-5</sup> | 4.8x10<sup>-3</sup> | 1.15 (1.06, 1.24) | 3.7x10<sup>-4</sup> | 0.026 |
| Uridine | 0.82 (0.75, 0.88) | 5.4x10<sup>-7</sup> | 1.5x10<sup>-5</sup> | 0.86 (0.79, 0.93) | 1.3x10<sup>-4</sup> | 0.010 | 0.86 (0.79, 0.93) | 1.7x10<sup>-4</sup> | 0.026 |
| 1-docosahexaenoylglycerophosphocholine | 0.82 (0.75, 0.90) | 2.2x10<sup>-5</sup> | 2.5x10<sup>-4</sup> | 0.85 (0.77, 0.93) | 3.6x10<sup>-4</sup> | 0.018 | 0.85 (0.77, 0.93) | 4.0x10<sup>-4</sup> | 0.026 |
| O-sulfo-L-tyrosine | 1.18 (1.09, 1.28) | 5.4x10<sup>-5</sup> | 5.5x10<sup>-4</sup> | 1.16 (1.07, 1.25) | 4.0x10<sup>-4</sup> | 0.018 | 1.12 (1.03, 1.23) | 0.01 | 0.170 |
| Glycoursodeoxycholate | 1.15 (1.08, 1.23) | 3.0x10<sup>-5</sup> | 3.4x10<sup>-4</sup> | 1.13 (1.05, 1.20) | 5.2x10<sup>-4</sup> | 0.018 | 1.13 (1.05, 1.20) | 5.4x10<sup>-4</sup> | 0.026 |
| Glycochenodeoxycholate | 1.16 (1.08, 1.24) | 1.8x10<sup>-5</sup> | 2.3x10<sup>-4</sup> | 1.13 (1.05, 1.21) | 5.8x10<sup>-4</sup> | 0.018 | 1.13 (1.06, 1.21) | 4.8x10<sup>-4</sup> | 0.026 |
| N-acetylalanine | 1.22 (1.14, 1.32) | 5.6x10<sup>-8</sup> | 2.7x10<sup>-6</sup> | 1.14 (1.06, 1.23) | 6.0x10<sup>-4</sup> | 0.018 | 1.11 (1.02, 1.21) | 0.02 | 0.205 |
| N-acetylthreonine | 1.21 (1.12, 1.31) | 7.3x10<sup>-7</sup> | 1.6x10<sup>-5</sup> | 1.14 (1.05, 1.23) | 9.2x10<sup>-4</sup> | 0.025 | 1.11 (1.02, 1.21) | 0.02 | 0.205 |
| Gamma-glutamylisoleucine | 0.86 (0.79, 0.94) | 4.6x10<sup>-4</sup> | 3.0x10<sup>-3</sup> | 0.87 (0.80, 0.95) | 1.5x10<sup>-3</sup> | 0.036 | 0.87 (0.80, 0.94) | 8.1x10<sup>-4</sup> | 0.031 |
| Gamma-glutamylleucine | 0.84 (0.78, 0.92) | 5.8x10<sup>-5</sup> | 5.7x10<sup>-4</sup> | 0.88 (0.81, 0.95) | 1.9x10<sup>-3</sup> | 0.042 | 0.87 (0.80, 0.94) | 1.0x10<sup>-3</sup> | 0.031 |
| FDR p: False Discovery Rate-adjusted p-values. Model 1: Proportional hazards model adjusted for age, sex, race, study site, and batch. Model 2: As Model 1, additionally adjusted for smoking, body mass index, systolic blood pressure, use of antihypertensive medication, diabetes mellitus, prevalent heart failure and prevalent coronary heart disease. Model 3: As Model 2, additionally adjusted for eGFR. |  |  |  |  |  |  |  |  |  |

**Figure 1.**
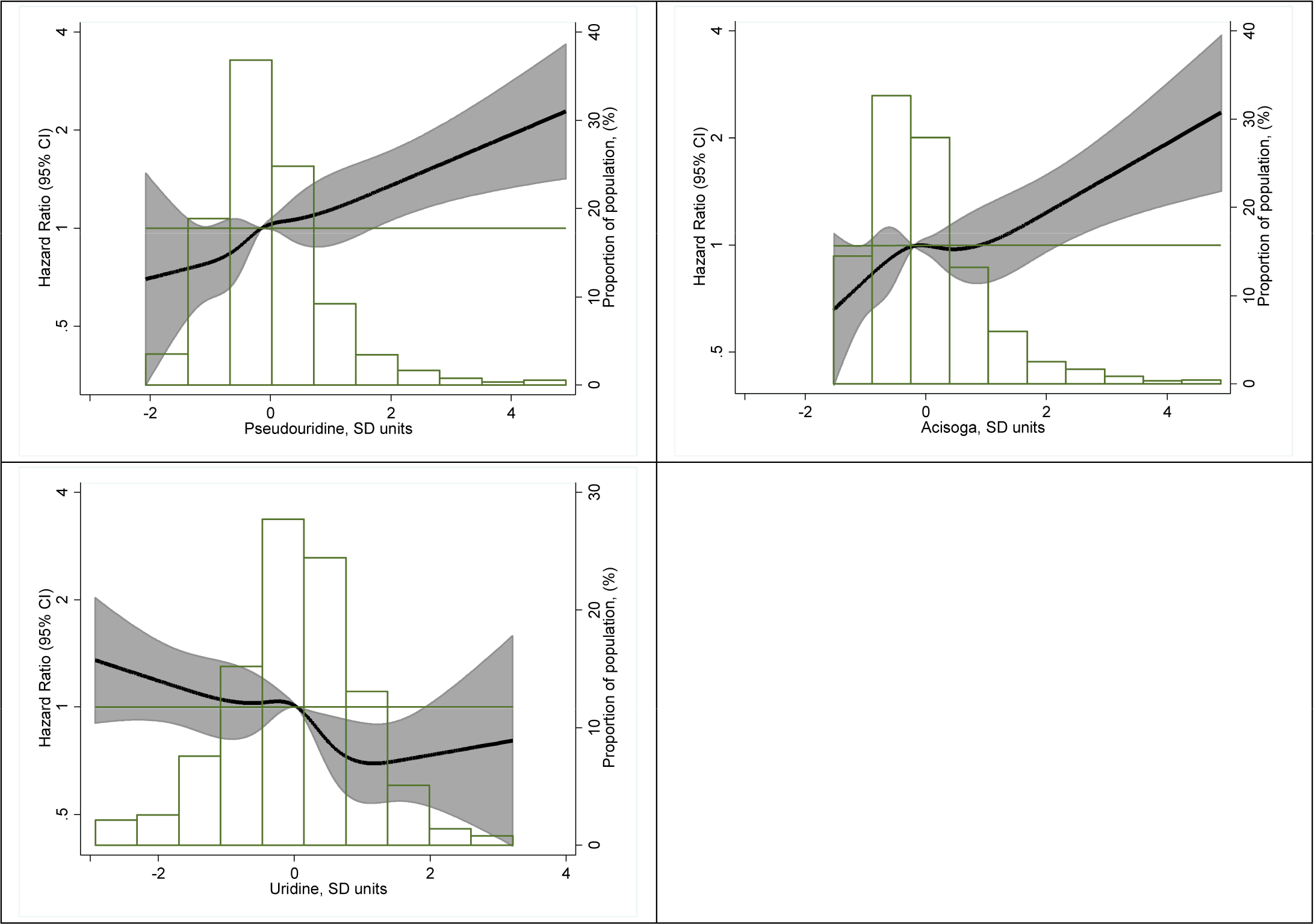
Association of concentrations of pseudouridine (top left panel), acisoga (top right panel) and uridine (bottom right panel) with incidence of atrial fibrillation presented as hazard ratio (HR; solid line) and 95% confidence intervals (CI; shaded area). Results from Cox proportional hazards model with metabolites modeled using restricted cubic splines (knots at 5th, 27.5th,50th, 72.5th, and 95th percentiles), adjusted for age, sex, race, batch, study site, body mass index, smoking, diabetes, systolic blood pressure, use of antihypertensive medication, prevalent coronary heart disease, and prevalent heart failure. Median value of the metabolite was considered the reference (HR = 1). The histograms represent the frequency distribution of metabolites levels. ARIC study, 1987–2013

To characterize in more detail the association of the three metabolites with AF, we explored their cross-sectional association with selected intermediate phenotypes of AF (PR interval, elevated P wave terminal force in V1, abnormal P wave axis) (Table 4). None of the three metabolites were associated with the odds of abnormal P wave axis or elevated P wave terminal force in V1. The results were suggestive of a possible association of higher pseudouridine and acisoga with shorter PR interval [beta (95% CI), −0.9 ms (−1.9, 0.1), and −0.9 ms (−1.8, −0.1), respectively) and higher uridine with longer PR interval [0.6 ms (−0.2, 1.4)].

**Table 4.** Association of pseudouridine, acisoga and uridine with selected ECG measures, ARIC study, 1987-1989

|  |  | PR duration, ms<sup>a</sup> |  | Abnormal P wave axis |  | Elevated P wave terminal force in V1 |  |
| --- | --- | --- | --- | --- | --- | --- | --- |
|  |  | Diff (95%CI) | P-value | OR (95%CI) | P-value | OR (95%CI) | P-value |
| Pseudouridine | Model 1 | 0.15 (-0.66, 0.97) | 0.71 | 0.86 (0.74, 1.00) | 0.04 | 1.11 (1.03, 1.20) | 0.01 |
|  | Model 2 | -0.56 (-1.39, 0.27) | 0.18 | 0.92 (0.79, 1.06) | 0.25 | 1.03 (0.94, 1.12) | 0.55 |
|  | Model 3 | -0.90 (-1.87, 0.06) | 0.07 | 0.91 (0.77, 1.08) | 0.27 | 1.00 (0.91, 1.11) | 0.92 |
| Acisoga | Model 1 | -0.52 (-1.31, 0.27) | 0.19 | 1.04 (0.92, 1.18) | 0.54 | 1.07 (0.99, 1.16) | 0.09 |
|  | Model 2 | -0.81 (-1.60, -0.02) | 0.05 | 1.01 (0.89, 1.15) | 0.86 | 1.03 (0.95, 1.11) | 0.51 |
|  | Model 3 | -0.90 (-1.71, -0.09) | 0.03 | 1.02 (0.89, 1.16) | 0.79 | 1.02 (0.94, 1.11) | 0.68 |
| Uridine | Model 1 | 0.79 (0.01, 1.57) | 0.05 | 0.92 (0.81, 1.04) | 0.17 | 0.96 (0.88, 1.04) | 0.30 |
|  | Model 2 | 0.58 (-0.21, 1.37) | 0.15 | 1.00 (0.88, 1.15) | 0.95 | 1.00 (0.92, 1.08) | 0.93 |
|  | Model 3 | 0.59 (-0.20, 1.38) | 0.15 | 1.00 (0.88, 1.15) | 0.96 | 1.00 (0.92, 1.09) | 0.98 |

Model 1: Adjusted for age, sex, race, study site, and batch. Model 2: As Model 1, additionally adjusted for smoking, body mass index, systolic
blood pressure, use of antihypertensive medication, diabetes mellitus, prevalent heart failure and prevalent coronary heart disease. Model 3: As
Model 2, additionally adjusted for eGFR
<sup>a</sup> Models additionally adjusted for resting heart rate

We assessed whether any of the AF-related genetic variants identified in a previously published GWAS of AF among individuals of European ancestry were associated with levels of pseudouridine, acisoga or uridine among white participants with genomic data (N = 1421). In this analysis, neither the individual genetic variants nor the AFGen genetic risk score predicted serum levels of these three metabolites (Supplementary Table 4).

Finally, variation in rs2272996 in gene *VNN1*, previously associated with circulating levels of acisoga, was not predictive of AF risk (p = 0.88 in the most recent GWAS from the AFGen consortium).

## DISCUSSION

In this metabolomic study of 3,922 men and women from a diverse prospective cohort we replicated a previously described association of glycocholenate sulfate, a secondary bile acid, with the incidence of AF. Also, we identified three additional metabolites (two related to pyrimidine metabolism, pseudouridine and uridine, and one related to polyamine metabolism, acisoga) associated with incidence of AF using a stringent Bonferroni correction. Several additional analyses showing lack of association of these metabolites with AF electrical endophenotypes and gene variants associated with AF in a previously published GWAS suggest that these metabolites may affect AF pathogenesis through alternative mechanisms. An additional analysis using FDR-adjusted significance thresholds identified eight additional metabolites associated with AF incidence, including two bile acids, two amino-acids, two dipeptides, one lisolipid, and one xenobiotic.

### Bile acids and AF

Consistent with our prior analysis of the ARIC cohort,^3^ we found an association of circulating glychocholenate sulfate with increased incidence of AF. The previously described association of another secondary bile acid, glycholithocholate sulfate, with AF was not replicated in this new analysis. In addition, we identified two additional secondary bile acids, glycoursodeoxycholate and glycochenodeoxycholate, associated with AF incidence. These associations did not achieve the multiple comparison-corrected threshold for statistical significance, but were significant using an FDR-adjusted p-value. Glychocholenate sulfate is possibly derived from 3-beta-hydroxy-5-cholenoic acid (cholenate). Prior literature has described elevations of cholenate in patients with liver disease,^16^ while both glycoursodeoxycholate and glycochenodeoxycholate are elevated in patients with liver cirrhosis.^17^ Thus, liver injury, which has been associated with AF previously, could explain the association of bile acids with incident AF. However, adjustment for biomarkers of liver damage (ALT and AST) did not materially change the associations. Alternative mechanisms, including the cardiometabolic implications of systemic activation of farnesoid X receptor by circulating bile acids^18^ or changes in the gut microbiota,^19^ instrumental in bile acid metabolism, could underlie the described associations. Our results, together with a prior study describing potential arrhythmogenic effects of bile acids,^20^ provide the rationale for future work exploring the impact of bile acids on the development of AF.

### Pseudouridine and uridine

Pseudouridine and uridine are nucleosides involved in RNA synthesis and metabolism. Pseudouridine results from enzymatic posttranscriptional modification of uridine in RNA, with stress conditions influencing the occurrence of this process.^21^ In turn, RNA pseudouridylation can affect gene expression regulation through mRNA stability and proteome diversity.^22^ Because of its physiological roles, circulating or urinary pseudouridine is considered a marker of RNA degradation and cell turnover.^23^ Prior studies have reported higher concentrations of circulating pseudouridine in patients with pulmonary arterial hypertension,^24^ heart failure,^25^ impaired kidney function,^26,27^ end-stage renal disease,^28,29^ and cancer.^30^ The relationships between circulating pseudouridine and posttranscriptional pseudouridylation of RNA and what role, if any, pseudouridine has in processes contributing to AF risk, requires further investigation.

Uridine is a ribonucleoside potentially involved in modulation of the metabolism of multiple systems and critical for cellular function and survival, though its specific targets have not been identified.^31^ Recent studies indicate that plasma uridine plays a key role in energy homeostasis and thermoregulation, modulating leptin signaling and potentially affecting glucose and insulin metabolism.^32^ Given the involvement of obesity and diabetes in the development of AF, deeper understanding of the physiological role of uridine in cardiometabolic disorders is needed. In fact, prior epidemiologic and clinical evidence has shown beneficial associations with higher plasma uridine, with higher levels of uridine associated with reduced mortality in the ARIC cohort,^33^ and reduced pulse wave velocity in the Twins UK Registry.^34^ In the Framingham Heart Study, higher concentrations of uridine were associated with a nonsignificant lower risk of AF (HR 0.84, 95%CI 0.70, 1.00, p = 0.05, per 1-standard deviation higher concentrations).^4^

### Acisoga

Acisoga (N-(3-acetamidopropyl)pyrrolidin-2-one) is a catabolic product of spermidine formed from N1-acetylspermidine, and involved in the metabolism of polyamines.^35^ Its precise role is unknown, but two prior studies have found associations of elevated acisoga concentrations with higher body mass index,^36,37^ and a potential association with the incidence of diabetes mellitus in the ARIC study.^38^ Concentrations of acisoga were part of a metabolomic-score predicting mortality in the Alpha-Tocopherol, Beta-Carotene Cancer Prevention study cohort.^39^ Polyamines are key players in a range of processes, including cell-cell interactions, cellular signaling, and ion channel regulation.^40^ Acisoga, as an end product of polyamine metabolism, may be a marker of dysregulation in this pathway.

### Other metabolites

Using a less stringent approach to define statistical significance, we identified additional metabolites associated with increased risk of AF. These included two amino acids (N-acetylalanine and N-acetylthreonine), two dipeptides (gamma-glutamylisoleucine and gamma-glutamylleucine), one lysolipid (1-docosahexaenoylglycerophosphocholine), and one xenobiotic (O-sulfo-L-tyrosine). To our knowledge, these metabolites have not been previously associated with the incidence of AF, other arrhythmias, or cardiovascular disease in general. Higher levels of N-acetylalanine and lower levels of gamma-glutamylleucine in blood have been associated with increased mortality in the ARIC cohort.^33^ In patients with type 1 diabetes, higher circulating levels of N-acetylalanine, N-acetylthreonine, and O-sulfo-L-tyrosine were associated with faster progression to end-stage renal disease,^29^ while N-acetylthreonine was associated with faster decline in kidney function in type 2 diabetes^27^ and N-acetylalanine and O-sulfo-L-tyrosine associated with incident chronic kidney disease in the general population.^26^ In our analysis, adjustment for kidney function attenuated the associations of N-acetylalanine, N-acetylthreonine, and O-sulfo-L-tyrosine with AF incidence, indicating they may be markers of impaired kidney function, an established risk factor for AF.^41^ Animal studies suggest that gamma-glutamylleucine is an indicator of anti-obesogenic metabolism,^42^ with obesity being a strong risk factor for AF.^43^ Decreased concentrations of lysolipids, including 1-docosahexaenoylglycerophosphocholine, have been associated with obesity in children.^36^

### Implications for understanding of AF pathophysiology

The circulating metabolites prospectively associated with the incidence of AF in the ARIC cohort highlight two major pathways in AF pathophysiology. First, several metabolites, including the amino acids N-acetylalanine and N-acetylthreonine, the xenobiotic O-sulfo-L-tyrosine and, to a lesser extent, pseudouridine, underline the role that kidney dysfunction plays in the development of AF.^41^ In addition to causing hypertension, chronic kidney disease can lead to chronic inflammation, abnormalities in calcium and phosphate metabolism, vascular dysfunction, and left ventricular hypertrophy, all potentially involved in AF development.^44^ A second pathway involves metabolic alterations associated with obesity, glycemic metabolism, and changes in the microbiome. The association of AF incidence with concentrations of secondary bile acids, acisoga, uridine, the lysolipid 1-docosahexaenoylglycerophosphocholine, and the dipeptides gamma-glutamylisoleucine and gamma-glutamylleucine fit within this pathway. Diabetes and obesity are established risk factors for AF,^45,46^ but the exact mechanisms responsible for this association are unclear. Our findings point to potential fruitful areas of further inquiry.

### Strengths and limitations

Our study has important strengths, including the inclusion of a large and diverse cohort with excellent follow-up, an adequate number of AF cases to identify associations, and the availability of extensive covariates to reduce confounding. Moreover, we have considered only metabolites that passed rigorous quality control criteria. However, the method of AF ascertainment—relying predominantly on hospital discharge diagnoses—has probably led to missed events, including asymptomatic AF and AF managed exclusively in outpatient settings. Other limitations include the risk of false negatives, due to the limited number of events, and the absence of an independent sample for replication.

### Future directions

Our findings identify potential fruitful avenues of research. Additional studies that aim to evaluate the role played by the metabolism of bile acids, uridine and polyamines in processes leading to AF are warranted. Replicating findings from the ARIC cohort in independent samples is also needed. Combining metabolomic data with those coming from other omic levels (genomics, transcriptomics, and proteomics) and exploring associations with intermediate phenotypes of AF (e.g. left atrial abnormalities) could be particularly rewarding.

### Conclusions

This study replicated the association of one bile acid with AF reported in a previous study and identified three additional metabolites from two metabolic pathways associated with AF. Our findings suggest that metabolomic approaches in large epidemiologic studies can be valuable in biomarker discovery and advancing our understanding of the pathogenesis of complex diseases.

## Supporting information

Supplemental Results

Complete results

## ACKNOWLEDGEMENTS

The authors thank the staff and participants of the ARIC study for their important contributions.

## AUTHORS’ CONTRIBUTIONS

AA, BY, and EB developed the initial study, with input from YVS, LYC, LRL, WTO and EZS. AA, LYC, LRL and ESZ contributed to obtain the information on atrial fibrillation incidence, while EB and BY led the acquisition of metabolomic data. AA, BY, YVS, LYC, and EB planned the analyses. BY and AA conducted statistical analysis. EB was responsible for study design, obtaining funding and providing supervision. AA wrote the initial draft of the manuscript. BY, YVS, LYC, LRL, WTO, EZS and EB read, provided critical comments, and approved the manuscript.

## DATA AVAILABILITY

Some access restrictions apply to the data underlying the findings. The consent signed by study participants does not allow the public release of their data. Data from the Atherosclerosis Risk in Communities Study can be accessed through the NHLBI BioLINCC repository (https://biolincc.nhlbi.nih.gov/home/) or by contacting the ARIC Coordinating Center (http://www2.cscc.unc.edu/aric/distribution-agreements).

