## Supplemental Results for "Serum metabolomics and incidence of atrial fibrillation: the Atherosclerosis Risk in Communities (ARIC) Study"

[1]Department of Epidemiology, Rollins School of Public Health, Emory University, Atlanta, GA. [2]University of Texas Health Sciences Center, Houston, TX. [3]Division of Cardiology, University of Minnesota Medical School, Minneapolis, MN. [4]University of North Carolina, Chapel Hill, NC. [5]Division of Cardiology, School of Medicine, Emory University, Atlanta, GA. [6]Department of Epidemiology and Prevention, Wake Forest School of Medicine, Winston-Salem, NC. [7]Human Genome Sequence Center, Baylor College of Medicine, Houston, TX

**Supplementary Table 1.** Pearson correlation coefficients between measurements obtained in 2010 and 2014 in paired serum baseline samples from 97 participants, ARIC study

| **Metabolite** | **Correlation coefficient** | **Missing, %** | |
|---|---|---|---|
| | | **Batch 1** | **Batch 2** |
| Pseudouridine | 0.74 | 0 | 0 |
| Acisoga | 0.57 | 0.5 | 0 |
| Uridine | 0.65 | 0 | 0.6 |
| 1-docosahexaenoylglycerophosphocholine | 0.46 | 0 | 0 |
| O-sulfo-L-tyrosine | 0.82 | 0.1 | <0.1 |
| Glycoursodeoxycholate | 0.97 | 16.1 | 0.4 |
| Glycochenodeoxycholate | 0.99 | 3.9 | 22.8 |
| N-acetylalanine | 0.71 | 0.1 | <0.1 |
| N-acetylthreonine | 0.41 | 3.0 | 0.2 |
| Gamma-glutamylisoleucine | 0.81 | 0.2 | 0 |
| Gamma-glutamylleucine | 0.77 | 0 | 0 |

**Supplementary Table 2.** Partial Pearson correlation between 11 top metabolites adjusted for batch effect.

| | Pseudouridine | N-acetylalanine | N-acetylthreonine | O-sulfo-L-tyrosine | Acisoga | GPCho 22:6n3 | Glycoursodeoxycholate | Glycochenodeoxycholate | gamma-glutamylisoleucine | gamma-glutamylleucine | Uridine |
|---|---|---|---|---|---|---|---|---|---|---|---|
| Pseudouridine | 1.00 | 0.68 | 0.58 | 0.61 | 0.42 | -0.01 | 0.08 | 0.11 | 0.11 | 0.08 | -0.02 |
| N-acetylalanine | 0.68 | 1.00 | 0.69 | 0.57 | 0.33 | 0.03 | 0.05 | 0.07 | 0.16 | 0.13 | 0.06 |
| N-acetylthreonine | 0.58 | 0.69 | 1.00 | 0.55 | 0.29 | 0.01 | 0.05 | 0.09 | 0.12 | 0.11 | -0.05 |
| O-sulfo-L-tyrosine | 0.61 | 0.57 | 0.55 | 1.00 | 0.28 | 0.02 | 0.08 | 0.06 | 0.08 | 0.06 | -0.01 |
| Acisoga | 0.42 | 0.33 | 0.29 | 0.28 | 1.00 | 0.06 | 0.03 | 0.09 | 0.03 | 0.07 | -0.03 |
| GPCho 22:6n3 | -0.01 | 0.03 | 0.01 | 0.02 | 0.06 | 1.00 | -0.01 | -0.03 | 0.04 | 0.06 | 0.08 |
| Glycoursodeoxycholate | 0.08 | 0.05 | 0.05 | 0.08 | 0.03 | -0.01 | 1.00 | 0.58 | -0.02 | -0.06 | -0.06 |
| Glycochenodeoxycholate | 0.11 | 0.07 | 0.09 | 0.06 | 0.09 | -0.03 | 0.58 | 1.00 | -0.03 | -0.09 | -0.12 |
| gamma-glutamylisoleucine | 0.11 | 0.16 | 0.12 | 0.08 | 0.03 | 0.04 | -0.02 | -0.03 | 1.00 | 0.91 | 0.13 |
| gamma-glutamylleucine | 0.08 | 0.13 | 0.11 | 0.06 | 0.07 | 0.06 | -0.06 | -0.09 | 0.91 | 1.00 | 0.15 |
| Uridine | -0.02 | 0.06 | -0.05 | -0.01 | -0.03 | 0.08 | -0.06 | -0.12 | 0.13 | 0.15 | 1.00 |

GPCho 22:6n3: 1-docosahexaenoylglycerophosphocholine (22:6n3)

**Supplementary Table 3**. Association of selected individual metabolites with incidence of atrial fibrillation after adjustment for blood lipids or exclusion of participants with prevalent cardiovascular disease at baseline, ARIC study, 1987-2013.

| Metabolite | Adjustment for blood lipids[a] (N = 3,873; AF = 606) | | Excluding participants with prevalent HF or CHD[b] (N = 3,567; AF = 514) | |
|---|---|---|---|---|
| | HR (95%CI) | P-value | HR (95%CI) | P-value |
| Pseudouridine | 1.15 (1.06, 1.26) | 0.002 | 1.19 (1.08, 1.31) | 0.0006 |
| Acisoga | 1.14 (1.05, 1.23) | 0.001 | 1.10 (1.01, 1.20) | 0.03 |
| Uridine | 0.87 (0.81, 0.94) | 0.0007 | 0.86 (0.79, 0.94) | 0.0007 |

[a] Model adjusted for age, sex, race, study site, batch, smoking, body mass index, systolic blood pressure, use of antihypertensive medication, diabetes mellitus, prevalent heart failure, prevalent coronary heart disease, eGFR, total serum cholesterol, total HDL cholesterol, and total triglycerides.

[b] Model adjusted for age, sex, race, study site, batch, smoking, body mass index, systolic blood pressure, use of antihypertensive medication, diabetes mellitus, prevalent heart failure, prevalent coronary heart disease, and eGFR.

**Supplementary Table 4**. Association of AFGen score and SNPs with metabolites in white participants of the ARIC study (N = 1481), 1987-89. Results from multiple linear regression models adjusted for age, sex and study site.

| | Pseudouridine | | Acisoga | | Uridine | |
|---|---|---|---|---|---|---|
| | Beta (95%CI) | P-value | Beta (95%CI) | P-value | Beta (95%CI) | P-value |
| AFGen score | -0.011 (-0.148, 0.126) | 0.87 | -0.013 (-0.152, 0.125) | 0.85 | 0.041 (-0.101, 0.184) | 0.57 |
| rs11264280 | -0.016 (-0.091, 0.060) | 0.68 | -0.024 (-0.100, 0.052) | 0.54 | -0.037 (-0.116, 0.041) | 0.35 |
| rs72700118 | -0.082 (-0.185, 0.021) | 0.12 | -0.101 (-0.205, 0.003) | 0.06 | -0.016 (-0.123, 0.090) | 0.76 |
| rs520525 | 0.039 (-0.038, 0.116) | 0.32 | 0.053 (-0.024, 0.131) | 0.18 | 0.033 (-0.047, 0.113) | 0.42 |
| rs2540949 | -0.001 (-0.072, 0.069) | 0.97 | -0.032 (-0.103, 0.039) | 0.38 | -0.007 (-0.080, 0.066) | 0.86 |
| rs3771537 | 0.023 (-0.045, 0.091) | 0.51 | 0.010 (-0.059, 0.079) | 0.77 | 0.014 (-0.057, 0.085) | 0.70 |
| rs2288327 | -0.022 (-0.113, 0.070) | 0.65 | -0.035 (-0.128, 0.058) | 0.46 | 0.064 (-0.031, 0.160) | 0.19 |
| rs11718898 | 0.069 (-0.005, 0.143) | 0.07 | 0.042 (-0.33, 0.117) | 0.27 | -0.007 (-0.084, 0.070) | 0.86 |
| rs6843082 | -0.025 (-0.109, 0.060) | 0.57 | 0.027 (-0.058, 0.113) | 0.53 | 0.011 (-0.076, 0.099) | 0.80 |
| rs337711 | -0.014 (-0.083, 0.055) | 0.69 | -0.014 (-0.083, 0.056) | 0.70 | 0.000 (-0.071, 0.072) | 0.99 |
| rs2967791 | 0.008 (-0.060, 0.076) | 0.82 | -0.031 (-0.100, 0.038) | 0.38 | -0.030 (-0.102, 0.041) | 0.40 |
| rs4946333 | -0.008 (-0.077, 0.060) | 0.81 | -0.027 (-0.097, 0.042) | 0.44 | -0.005 (-0.077, 0.066) | 0.88 |
| rs12664873 | 0.029 (-0.045, 0.102) | 0.44 | -0.025 (-0.099, 0.050) | 0.52 | -0.055 (-0.131, 0.022) | 0.16 |
| rs1997572 | 0.026 (-0.046, 0.097) | 0.48 | 0.041 (-0.032, 0.113) | 0.27 | 0.015 (-0.059, 0.090) | 0.69 |
| rs7508 | 0.048 (-0.029, 0.125) | 0.23 | 0.004 (-0.074, 0.082) | 0.92 | -0.021 (-0.101, 0.059) | 0.61 |
| rs7026071 | -0.012 (-0.083, 0.058) | 0.73 | 0.023 (-0.049, 0.094) | 0.53 | 0.043 (-0.030, 0.117) | 0.25 |
| rs7915134 | 0.035 (-0.061, 0.131) | 0.48 | 0.018 (-0.079, 0.115) | 0.71 | 0.108 (0.009, 0.208) | 0.03 |
| rs11598047 | 0.014 (-0.080, 0.107) | 0.78 | -0.036 (-0.130, 0.059) | 0.46 | 0.030 (-0.067, 0.127) | 0.54 |

| | | | | | | |
|---|---|---|---|---|---|---|
| rs35176054 | -0.036 (-0.135, 0.064) | 0.48 | 0.099 (-0.001, 0.200) | 0.05 | 0.029 (-0.074, 0.132) | 0.58 |
| rs75190942 | 0.121 (-0.033, 0.274) | 0.12 | -0.105 (-0.261, 0.050) | 0.18 | -0.075 (-0.234, 0.085) | 0.36 |
| rs883079 | 0.016 (-0.059, 0.090) | 0.68 | -0.012 (-0.087, 0.063) | 0.76 | 0.005 (-0.072, 0.082) | 0.90 |
| rs1152591 | -0.067 (-0.137, 0.002) | 0.06 | -0.048 (-0.118, 0.023) | 0.18 | -0.015 (-0.087, 0.058) | 0.69 |
| rs74022964 | -0.037 (-0.134, 0.059) | 0.45 | 0.044 (-0.054, 0.141) | 0.38 | 0.029 (-0.072, 0.129) | 0.57 |
| rs2106261 | -0.031 (-0.123, 0.061) | 0.50 | -0.035 (-0.127, 0.058) | 0.47 | -0.010 (-0.105, 0.086) | 0.84 |

**Supplementary Figure 1.** Hazard ratios (HR) and 95% confidence intervals (95%CI) of atrial fibrillation by 1-standard deviation difference in levels of glycholenate sulfate (top panel) or pseudouridine (bottom panel), adding individual covariates to Model 1 (including age, sex, center, race and batch, if applicable). ARIC study, 1987-2013. BMI: body mass index; BP: blood pressure; CHD: coronary heart disease; HF: heart failure. Model 2 includes covariates in Model 1 and all other covariates in the figure.

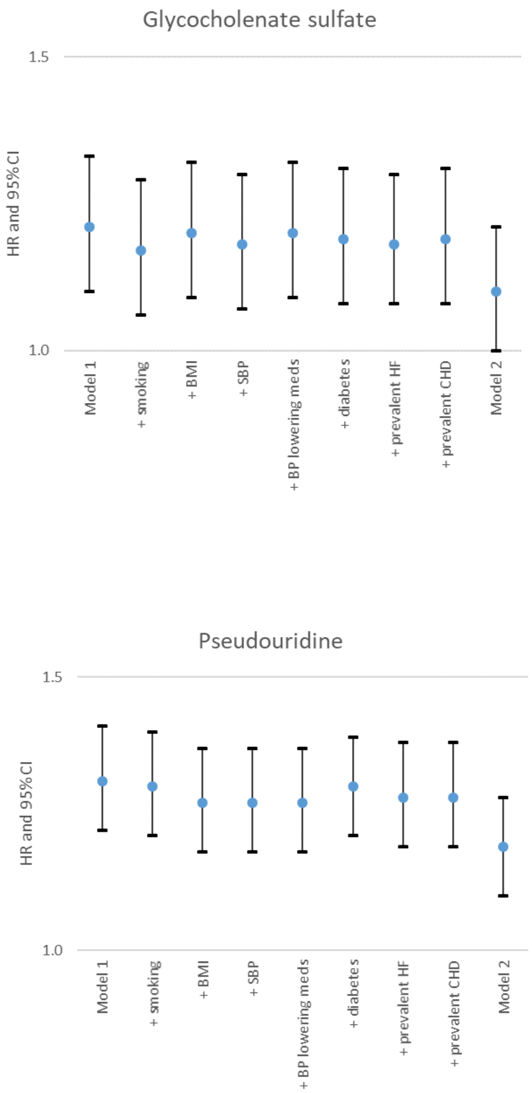

**Supplementary Figure 2**. Association of selected metabolites with incidence of atrial fibrillation by race, ARIC study, 1987–2013. Cox proportional hazards model adjusted for age, sex, study site, batch, smoking, body mass index, systolic blood pressure, use of antihypertensive medication, diabetes

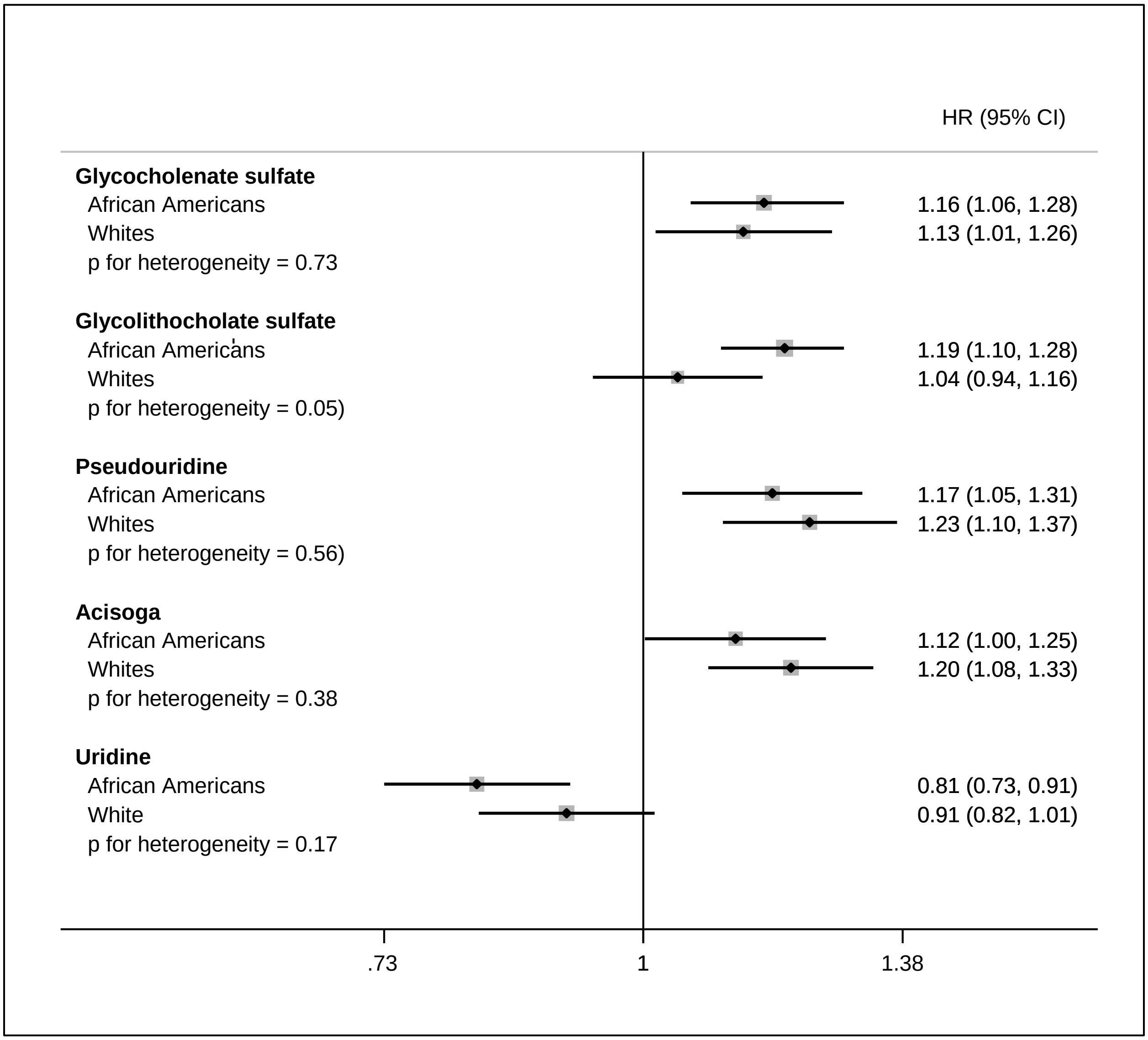


mellitus, prevalent heart failure and prevalent coronary heart disease.

**Supplementary Figure 3**. Association of selected metabolites with incidence of atrial fibrillation by sex, ARIC study, 1987–2013. Cox proportional hazards model adjusted for age, race, study site, batch, smoking, body mass index, systolic blood pressure, use of antihypertensive medication, diabetes mellitus, prevalent heart failure and prevalent coronary heart disease.

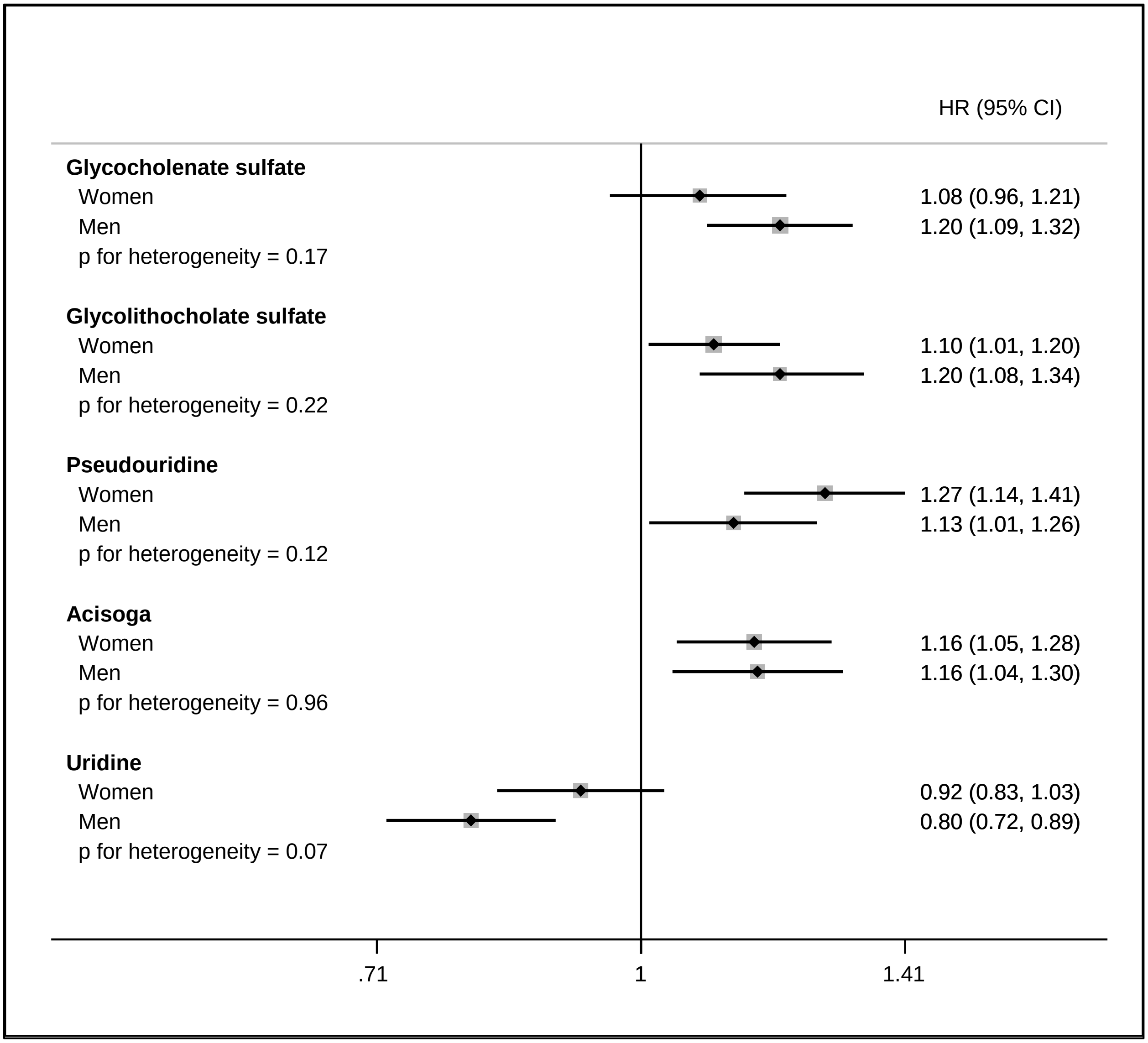
